# Molecular basis underlying specific interaction of mammalian melanopsins with an antagonist AA92593

**DOI:** 10.1101/2024.11.01.621458

**Authors:** Kohei Obayashi, Ruisi Zou, Toshifumi Mori, Hisao Tsukamoto

**Affiliations:** Department of Biology, Graduate School of Science, Kobe University, Kobe, Japan; Interdisciplinary Graduate School of Engineering Sciences, Kyushu University, Fukuoka, Japan; Institute for Materials Chemistry and Engineering, Kyushu University, Fukuoka, Japan; Center of Optical Scattering Image Science, Kobe University, Japan

## Abstract

Melanopsin functions in intrinsically photosensitive retinal ganglion cells of mammals to regulate circadian clock and pupil constriction. The opsinamide AA92593 has been reported to specifically inhibit mouse and human melanopsin functions as a competitive antagonist against retinal; however, the molecular mechanisms underlying its specificity have not been resolved. In this study, we attempted to identify amino acid residues responsible for the specific interaction of AA92593 with mammalian melanopsins. Our cell-based assays confirmed that AA92593 effectively inhibited the light-induced cellular responses of mammalian melanopsins, but not those of non-mammalian vertebrate and invertebrate melanopsins. These results suggest that amino acid residues specifically conserved among mammalian melanopsins are important for the antagonistic effect of AA92593, and we noticed Phe-94, Ser-188, and Ser-269 as candidate residues. Substitutions of these residues reduced the antagonistic effect of AA92593. We conducted docking and molecular dynamics simulations based on the AlphaFold-predicted melanopsin structure. The simulations indicated that Phe-94, Ser-188, and Ser-269 are located at the AA92593-binding site, and additionally identified Trp-189 and Leu-207 interacting with the antagonist. Substitutions of Trp-189 and Leu-207 affected the antagonistic effect of AA92593. Furthermore, substitutions of these amino acid residues converted AA92593-insensitive melanopsins susceptible to the antagonist. Based on experiments and molecular simulations, five amino acid residues, at positions 94, 188, 189, 207, and 269, were found to be responsible for the specific interaction with AA92593 in mammalian melanopsins.

## Introduction

In mammals, melanopsin (or Opn4) is expressed in intrinsically photosensitive retinal ganglion cells (ipRGCs) and plays important roles in non-visual photoreceptive functions such as circadian photoentrainment and pupil constriction (1, 2). Melanopsin is a blue light-sensitive G protein-coupled receptor (GPCR) and a member of the opsin family (3, 4). Light-induced regulation of melanopsin function has been utilized in neuroscience, behavioral, and clinical studies (5–7). Melanopsin functions can also be manipulated by chemicals. An opsinamide AA92593 was reported to be a competitive antagonist against the chromophore retinal, specifically acting on melanopsin (8). The original study clearly showed that AA92593 effectively antagonized the activity of melanopsin-expressing ipRGCs, but not visual photoreceptor cells. Since then, several studies have used AA92593 to suppress melanopsin function chemically (5, 9, 10). However, it is unclear how AA92593 selectively blocks the photoresponse of mammalian-type melanopsin (Opn4m) and whether an antagonistic effect is observed in closely related opsins such as non-mammalian-type melanopsin (Opn4x). To understand the specificity of AA92593 for mammalian melanopsins, it is necessary to identify the amino acid residues that are responsible for its antagonistic effect. Unfortunately, structural information about melanopsin is limited, despite the recent progress in GPCR structural biology (11).

In the present study, we identified the amino acid residues responsible for the antagonistic effect of AA92593. We performed site-directed mutagenesis and several cell-based assays on human and mouse melanopsins to evaluate the antagonist-induced effects on melanopsin mutants. We also conducted docking and molecular dynamics (MD) simulations of human melanopsin with AA92593, based on an AlphaFold-predicted structure, to visualize the interaction between melanopsin and the antagonist. Mutational experiments and computational simulations produced consistent results, indicating that five amino acid residues near the retinal-binding site in mammalian melanopsin directly interact with AA92593. Substitutions of the corresponding amino acid residues in AA92593-insensitive non-mammalian and invertebrate melanopsins increased their susceptibility to the antagonist. Our data revealed how AA92593 specifically acts as an antagonist against mammalian melanopsins, and the data provide valuable information for engineering melanopsins to be effectively (or ineffectively) regulated by the antagonist.

## Results

### Assessment of antagonistic effects of AA92593 on mammalian melanopsin (Opn4m) using GsX GloSensor assays

To quantify the antagonistic effects of AA92593 on mammalian melanopsins, we used the GsX GloSensor assay (12, 13). This assay measures GPCR-induced increases in intracellular cAMP levels even for Gq-, Gi/o-, or G12-coupled receptors by using Gsα chimeras (12, 13). Because melanopsin is primarily coupled with Gq-type G proteins, a Gsα mutant with 11 C-terminal sequence of Gqα (Gsα/q11) was expressed in COS-1 cells with melanopsin and a luciferase-based cAMP biosensor (see “Experimental Procedures”) (14). This assay enables the measurement of Gq activation by melanopsin as cAMP production (an increase in luminescence). In this study, we used C-terminal truncated melanopsin constructs to increase the expression levels with minimal compromise in functionality (9, 15–18). Previous studies of melanopsin have reported that the C-terminal truncation does not affect the photoreaction or G protein activation of melanopsins. In addition, the C-terminal region is located far from the binding site of AA92593 (see below). Hereafter, the C-terminal truncated melanopsin constructs are named as “WT”.

Our GsX GloSensor assay successfully detected Gsα/q11 activation by human and mouse melanopsins as a light-dependent increase in intercellular cAMP levels (Fig. 1, A and B). The addition of AA92593 effectively suppressed the light-dependent response in a concentration-dependent manner (Fig. 1, A and B), indicating that its antagonistic effect on the melanopsins can be detected using this assay. The inhibition of human melanopsin activity was dependent on the AA92593 concentration (Fig. 1C, EC_50_ = 1.05±0.28 μM). A similar antagonistic effect of AA92593 was observed for mouse melanopsin (Fig. 1D, EC_50_ = 2.98±0.58 μM). The lower EC_50_ value of human melanopsin compared with mouse melanopsin is likely due to its lower affinity for retinal (9).

**Fig. 1.**
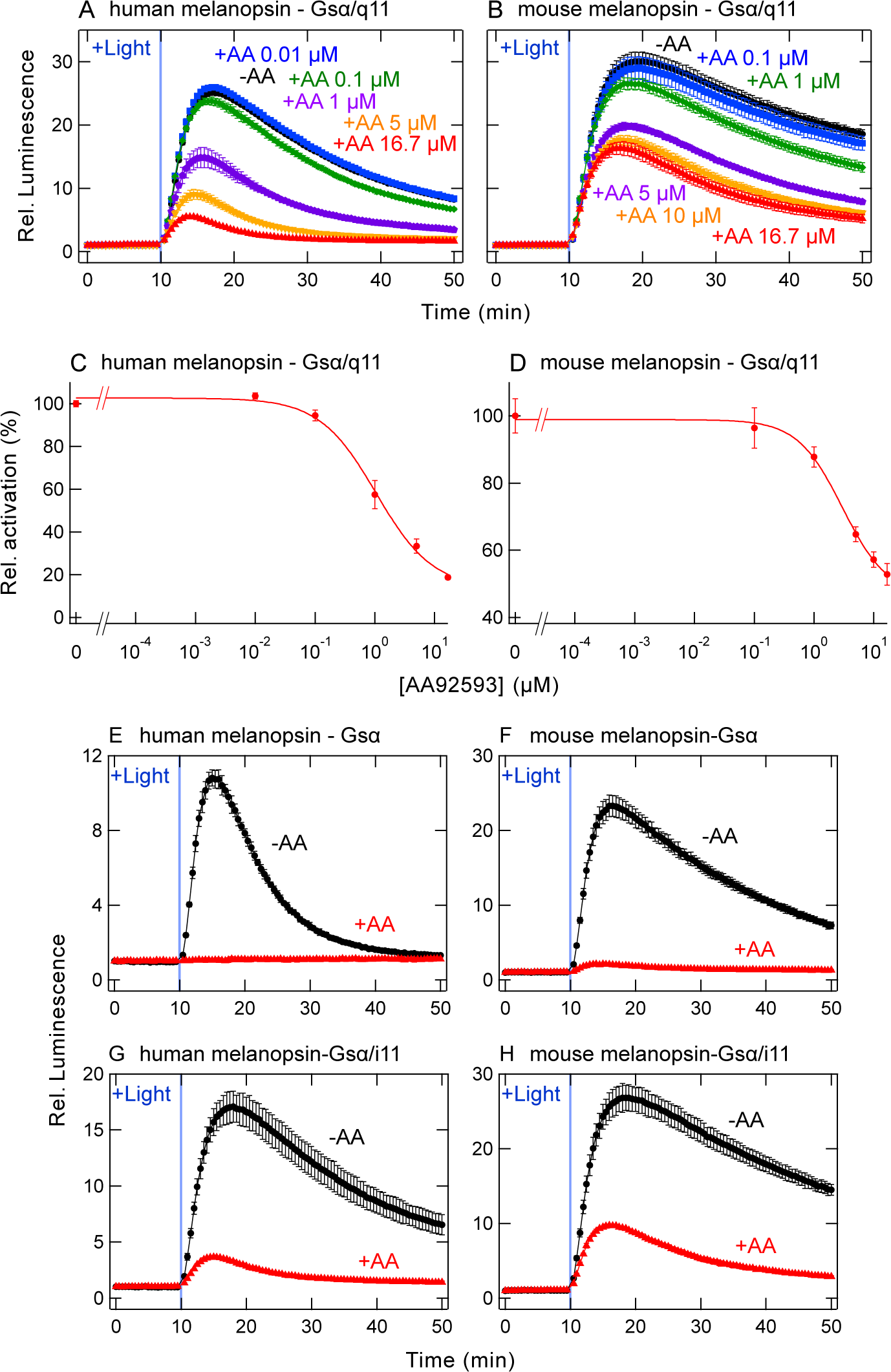
AA92593-induced suppression of G protein activation by mammalian melanopsins assessed by GsX Glosensor assay. (*A* and *B*) AA92593 concentration-dependent inhibition of intracellular cAMP elevation in COS-1 cells upon Gsα/q11 activation of human (*A*) and mouse (*B*) melanopsins. Luminescence levels of cAMP biosensor (GloSensor) are normalized to the values at the starting point (time = 0 min). In panel *A*, final AA92593 concentrations are 16.7 μM (red), 5 μM (orange), 1 μM (purple), 0.1 μM (green), 0.01 μM (blue), and 0 (black), respectively. In panel *B*, final AA92593 concentrations are 16.7 μM (red), 10 μM (orange), 5 μM (purple), 1 μM (green), 0.1 μM (blue), and 0 (black), respectively. Light blue bars indicate white light illumination (10 sec). Error bars indicate the SD values (n = 3). (*C* and *D*) Dose-dependent reduction in peak cAMP responses upon Gsα/q11 activation in human (*C*) and mouse (*D*) melanopsin-expressing COS-1 cells. Relative average peak values to the value in the absence of AA92593 are plotted against the final concentrations of AA92593. Error bars indicate the SD values (n = 3). EC_50_ values for-human and mouse melanopsins are 1.05±0.28 μM and 2.98±0.58 μM, respectively. (*E* and *F*) AA92593-dependent inhibition of intracellular cAMP elevation in COS-1 cells upon Gsα activation of human (*E*) and mouse (*F*) melanopsins. Error bars indicate the SD values (n = 3). (*G* and *H*) AA92593-dependent inhibition of intracellular cAMP elevation in COS-1 cells upon Gsα/i11 activation of human (*G*) and mouse (*H*) melanopsins. Error bars indicate the SD values (n = 3). In panels *E*-*H*, Red and black curves indicate luminescence changes in the presence and absence of 16.7 μM AA92593, respectively. Light blue bars indicate white light illumination (10 sec). Luminescence levels of cAMP biosensor (GloSensor) are normalized to the values at the starting point (time = 0 min).

Previous studies of human and mouse melanopsins reported that these melanopsins can activate not only Gq but Gs and Gi (19–21). Thus, we assessed the effect of AA92593 on Gsα WT or Gsα/i11 (with 11 C-terminal sequence of Giα) using the GsX GloSensor assay. The results indicated that AA92593 can suppress melanopsin-induced cAMP responses via exogenous Gsα (Fig. 1, E and F) or Gsα/i11 (Fig. 1, G and H). In addition, both mammalian melanopsins induced light-dependent cAMP responses via endogenous G proteins in COS-1 cells, and the endogenous G protein-induced cAMP responses were inhibited by the addition of AA92593. The endogenous G proteins produced much less luminescence signals in the GloSensor assay, probably due to the low expression levels of endogenous G proteins (Fig. S1). These results showed that the activation of Gq, Gs, and Gi by photo-activated melanopsins was suppressed by AA92593, which is consistent with the previous study showing that the molecule competes with the chromophore retinal for binding to melanopsin (8).

### Conserved amino acid residues important for susceptibility to AA92593

Because the inhibitory effect of AA92593 was detected by the GsX GloSensor assay, we attempted to identify the amino acid residues important for the susceptibility to AA92593. Because AA92593 competes with the retinal chromophore for binding to mammalian melanopsins (8), we noticed amino acid residues near the retinal-binding site that are specifically conserved among mammalian melanopsins. Based on sequence alignment of amino acid residues constituting the putative retinal-binding site (Fig. S2), three residues, Phe-94, Ser-188, and Ser-269, were found to be conserved in mammalian-type melanopsin (Opn4m) (Figs. 2A and S2). In non-mammalian-type melanopsin (Opn4x), these positions are not conserved, and are occupied by Cys, Thr, and Ala, respectively (Figs. 2A and S2).

**Fig. 2.**
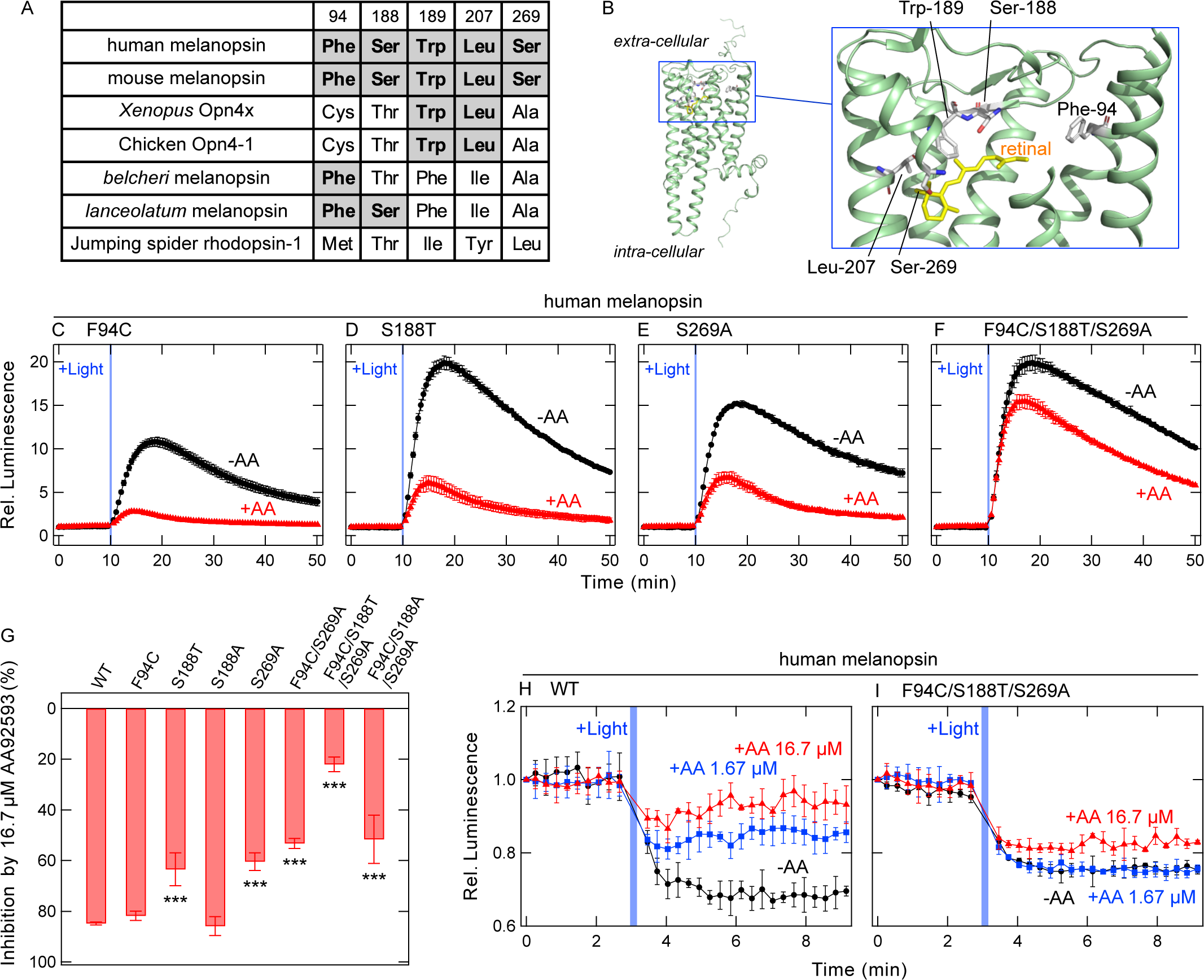
Changes in susceptibility of human melanopsin for AA92593 by substitutions of conserved amino acid residues in the putative retinal-binding site. (*A*) Amino acid residues at positions 94, 188, 189, 207, and 269 in various melanopsins and a non-melanopsin Gq-coupled opsin. “Mammalian melanopsin-type” amino acids are highlighted as bold. The amino acid residue numbering in this paper is based on the amino acid sequence of bovine rhodopsin. (*B*) Arrangement of the amino acid residues at positions 94, 188, 189, 207, and 269 in the AlphaFold2-predicted human melanopsin structure (51). The 11-*cis*-retinal molecule is adopted from the crystal structure of bovine rhodopsin (PDB ID: 1U19) (52). Note that the predicted structure of human melanopsin and the crystal structure of bovine rhodopsin with 11-*cis*-retinal were overlapped to obtain a structure that looks like retinal bound to human melanopsin. The structural models are prepared using PyMOL (https://pymol.org/). (*C*-*F*) AA92593-dependent inhibition of intracellular cAMP elevation in COS-1 cells upon Gsα/q11 activation of human melanopsin mutants F94C (*C*), S188T (*D*), S269A (*E*), and F94C/S188T/S269A (*F*). Red and black traces indicate luminescence changes in the presence and absence of 16.7 μM AA92593, respectively. Light blue bars indicate white light illumination (10 sec). Luminescence levels of cAMP biosensor (GloSensor) are normalized to the values at the starting point (time = 0 min). Error bars indicate the SD values (n = 3). (*G*) Comparison of inhibition in peak cAMP responses by 16.7 μM AA92593 upon Gsα/q11 activation in human melanopsin WT and mutants. Error bars indicate the SD values (n = 46, 8, 4, 3, 7, 6, 7, and 4 for WT, F94C, S188T, S188A, S269A, F94C/S269A, F94C/S188T/S269A and F94C/S188A/S269A, respectively). The statistical *P* values in differences from WT are 0.99, <0.001***, 1.00, <0.001***, <0.001***, <0.001***, and <0.001*** for F94C, S188T, S188A, S269A, F94C/S269A, F94C/S188T/S269A, and F94C/S188A/S269A, respectively (Dunnett’s test following one-way ANOVA). (*H* and *I*) NanoBiT Gq dissociation assay using Gqα/R183Q-LgBiT on human melanopsin WT (*H*) and F94C/S188T/S269A mutant (*I*). Red, blue, and black traces indicate luminescence changes in the presence of 16.7 μM (red), 1.67 μM (blue), and no (black) AA92593. Light blue bars indicate white light illumination (10 sec). NanoLuc luminescence levels are normalized to the values at the starting point (time = 0 min). Error bars indicate the SD values (n = 3).

We substituted the residues Phe-94, Ser-188, and Ser-269 in human melanopsin (Fig. 2, A and B) and examined the inhibitory effect of AA92593 on these mutants. Substitution of these amino acid residues with the corresponding amino acids in non-mammalian-type melanopsins (F94C, S188T, and S269A) reduced the inhibitory effect of AA92593 compared to its effect on WT (Fig. 2, C-E). While statistically significant difference was not observed between the F94C mutant and WT, the inhibitory effect was consistently weaker for the mutant in each experiment (Fig. 2, C and G, and Fig. S3). The antagonistic effect of AA92593 was significantly reduced against the S188T and S269A mutants (Fig. 2, D, E, and G). Unlike the S188T substitution, the S188A substitution did not induce a significant change in the effect of AA92593 (Fig. 2G). Similar results were observed for the mouse melanopsin F94C, S188T, and S269A mutants (Fig. 3, A, B, C, and E), although only the S269A mutant showed a statistically significant difference from WT (see below).

**Fig. 3.**
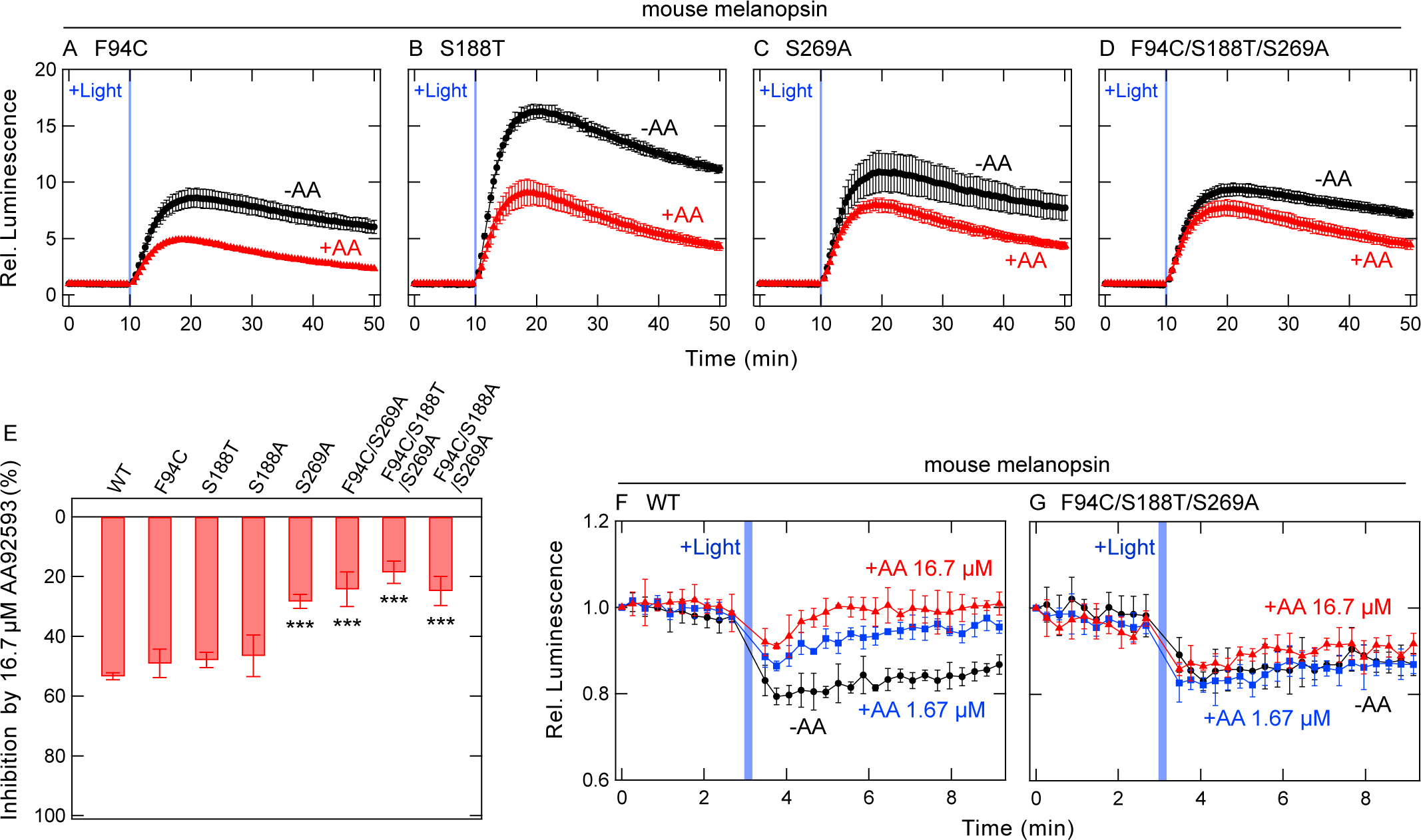
Changes in susceptibility of mouse melanopsin for AA92593 by substitutions in the putative retinal-binding site. (*A*-*D*) AA92593-dependent inhibition of intracellular cAMP elevation in COS-1 cells upon Gsα/q11 activation of mouse melanopsin mutants F94C (*A*), S188T (*B*), S269A (*C*), and F94C/S188T/S269A (*D*). Red and black traces indicate luminescence changes in the presence and absence of 16.7 μM AA92593, respectively. Light blue bars indicate white light illumination (10 sec). Luminescence levels of cAMP biosensor (GloSensor) are normalized to the values at the starting point (time = 0 min). Error bars indicate the SD values (n = 3). (E) Comparison of inhibition in peak cAMP responses by 16.7 μM AA92593 upon Gsα/q11 activation in mouse melanopsin WT and mutants. Error bars indicate the SD values (n = 23, 5, 4, 3, 5, 6, 6, and 5 for WT, F94C, S188T, S188A, S269A, F94C/S269A, F94C/S188T/S269A and F94C/S188A/S269A, respectively). The statistical *P* values in differences from WT are 0.95, 0.88, 0.81, <0.001***, <0.001***, <0.001***, and <0.001*** for F94C, S188T, S188A, S269A, F94C/S269A, F94C/S188T/S269A and F94C/S188A/S269A, respectively (Dunnett’s test following one-way ANOVA). (*F* and *G*) NanoBiT Gq dissociation assay on mouse melanopsin WT (*F*) and F94C/S188T/S269A mutant (*G*). Red, blue, and black traces indicate luminescence changes in the presence of 16.7 μM (red), 1.67 μM (blue), and no (black) AA92593. Light blue bars indicate white light illumination (10 sec). NanoLuc luminescence levels are normalized to the values at the starting point (time = 0 min). Error bars indicate the SD values (n = 3).

The single substitutions F94C, S188T, and S269A in mammalian melanopsins reduced the inhibitory effects of AA92593 (Figs. 2G and 3E). We expected combination of these substitutions to further decrease the AA92593 effect. We introduced the triple substitutions F94C/S188T/S269A into human and mouse melanopsins, and examined the inhibitory effects of AA92593. In both mammalian melanopsins, the F94C/S188T/S269A substitution reduced the inhibitory effect of AA92593 more than a single substitution (Figs. 2F, 2G, 3D, and 3E). These results clearly indicated that the effects of Phe-94, Ser-188, and Ser-269 on the susceptibility to AA92593 are additive in mammalian melanopsins.

As described above, we have assessed the inhibitory effect of AA92593 using the GsX GloSensor assay. To confirm whether an antagonistic effect could be observed in another experimental system, we used the NanoBiT G protein dissociation assay (22). In this assay, Gq activation (dissociation) is detected as a decrease in NanoLuc luminescence from the Lg-BiT fragment inserted with Gqα and the Sm-BiT fragment fused with Gβγ (see “Experimental Procedures”) (22, 23). If an antagonistic effect is observed in the NanoBiT assay, the light-dependent decrease in luminescence (Gq dissociation) by melanopsin would be reduced in the presence of AA92593. As expected, human melanopsin WT showed a light-dependent decrease in luminescence intensity (Fig. S4), indicating that Gq activation by the melanopsin was successively detected using the NanoBiT assay. A recent study reported that the introduction of a GTPase-deficient substitution, R183Q, on Gqα increased GPCR-induced Gq dissociation signals in a BRET-based biosensor assay, TRUPATH (24). Based on this insight, we prepared the Gqα with Lg-BiT containing the R183Q mutation and confirmed that melanopsin-induced Gq dissociation signals on NanoBiT assay was larger than Gqα WT with Lg-BiT (Fig. S4), probably because of the suppression of Gq inactivation in the Gqα R183Q mutant. We used the Gqα/R183Q-LgBiT to compare the signals induced by melanopsin mutants, with or without AA92593.

The light-dependent decrease in luminescence was weakened by the addition of AA92593 in a concentration-dependent manner (Fig. 2H). The human melanopsin mutant F94C/S188T/S269A also showed a robust light-dependent decrease in NanoLuc luminescence intensity. However, the AA92593-dependent inhibitory effect on Gq activation by the mutant was much reduced compared to that on WT (Fig. 2I). The antagonistic effect on Gq activation by mouse melanopsin was similarly reduced by the F94C/S188T/S269A substitutions (Fig. 3, F and G). These NanoBiT assay data clearly indicate that the antagonistic effect of AA92593 on mammalian melanopsins was observed at the Gq activation levels, which supports our conclusion that Phe-94, Ser-188, and Ser-269 play important roles in the susceptibility.

### Docking and MD simulations of human melanopsin with AA92593

To visualize how Phe-94, Ser-188, and Ser-269 make mammalian melanopsin susceptible to AA92593, we conducted computational studies on the human melanopsin-AA92593 complex. Despite recent progress in structural analyses of various GPCRs, the structure of melanopsin has not yet been experimentally solved. Thus, we adopted an AlphaFold-predicted structural model of human melanopsin and performed docking analysis with AA92593. Calculations using the widely used program AutoDock (25) were unsuccessful in obtaining the ligand bound to the pocket of human melanopsin, possibly because of steric clashes with the side-chains (26) (Fig. S5). In contrast, another program, DiffDock (27), obtained a complex structure in which the ligand was located in the binding pocket of the human melanopsin. The receptor-ligand docked structure with the best score clearly indicated that Ser-188 and Ser-269 were located in the binding site of AA92593, whereas Phe-94 was slightly away from AA92593 (Fig. 4A). These locations were consistent with our experiments, showing that single substitutions of S188T and S269A significantly reduced the antagonistic effect, whereas the single F94C substitution resulted in a minor reduction (Fig. 2G). In addition, Ala-114, Gly-117, Ala-118, Trp-189, Leu-207, Cys-208, Trp-265, Tyr-268, and Ala-272 were found to be located within 4 Å from the ligand (Fig. 4B).

**Fig. 4.**
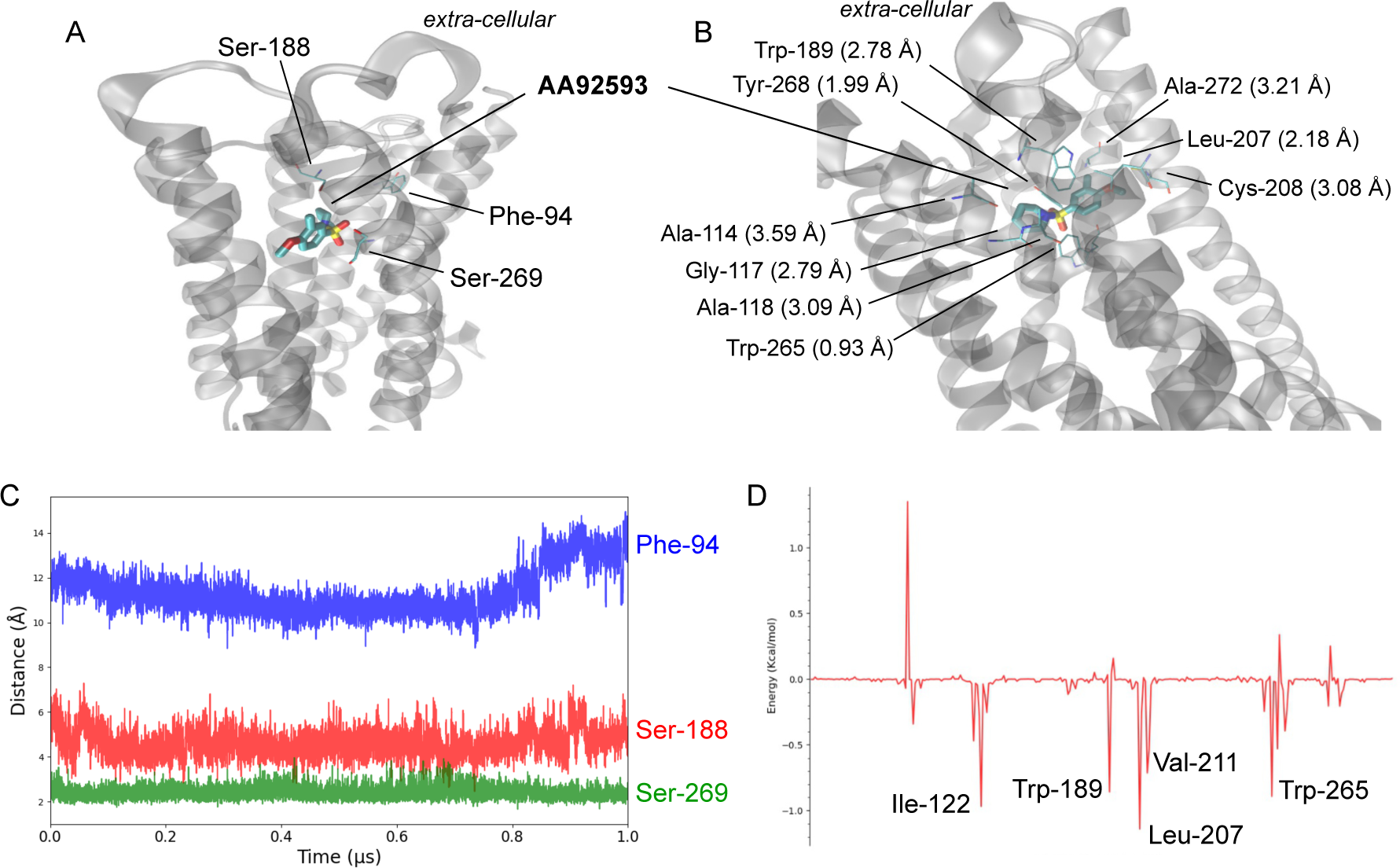
Docking and MD simulations on human melanopsin with AA92593. (*A*, *B*) Structure of the human melanopsin-AA92593 complex predicted from DiffDock. Parenthesis in panel B show the shortest distances between AA92593 and selected ligands. (*C*) Time evolution of the distances between AA92593 and Phe-94 (blue), Ser-188 (red), and Ser-269 (green) over the 1 µs MD trajectory. (*D*) Per-residue decomposition of the binding energy calculated using MMPBSA.

Since the ligand in the binding pocket predicted in crystal structures may not always be rigid at the binding pocket under the physiological conditions (28, 29), and the protein environment of the ligand may also show conformational changes over time (30, 31), we next explored the dynamics of the melanopsin-AA92593 interactions in the complex by performing MD simulations. The initial structure was obtained from the docking model, and the membrane and solvents were added as described below. The ligand remained in the binding pocket throughout the 1 µs-long simulation, while occasionally adjusting its orientation inside the pocket. In agreement with the docking model, Ser-188 and Ser-269 remained near the ligand while Phe-94 was mostly > 10 Å away from the ligand (Fig. 4C). The per-residue decomposition of the binding energy, obtained by the molecular mechanics Poisson-Boltzmann surface area (MMPBSA) calculations, indicated that Ile-122, Trp-189, Leu-207, Val-211, and Trp-265 contribute significantly to stabilizing binding, primarily via van der Waals interactions (Figs. 4D and S6). The distances between the ligand and these residues were mostly within 3 Å (Fig. S7). Notably, not all residues in contact with the ligand showed a large binding energy (e.g., Cys-208) (Fig. 4B), implying that both the distance and character of the residues were important for stabilization.

Docking and MD simulations of human melanopsin docked with AA92593 identified several amino acid residues that presumably interact with the antagonist in addition to Phe-94, Ser-188, and Ser-269. In particular, Trp-189, Leu-207, and Trp-265, which are conserved among not only mammalian-type but also non-mammalian-type melanopsins (Fig. S2), were found to be important for AA92593 binding in both the docking and MD results (Figs. 4, S6, and S7). Because Trp-265 is highly conserved in GPCRs and has been reported to be important for proper function and/or folding in many GPCRs, including opsins, the Trp-265 substitution tends to severely impair receptor functions (32–35). Thus, we assessed whether substitutions of Trp-189 and Leu-207 affect the susceptibility of human melanopsin to AA92593.

**Fig. 5.**
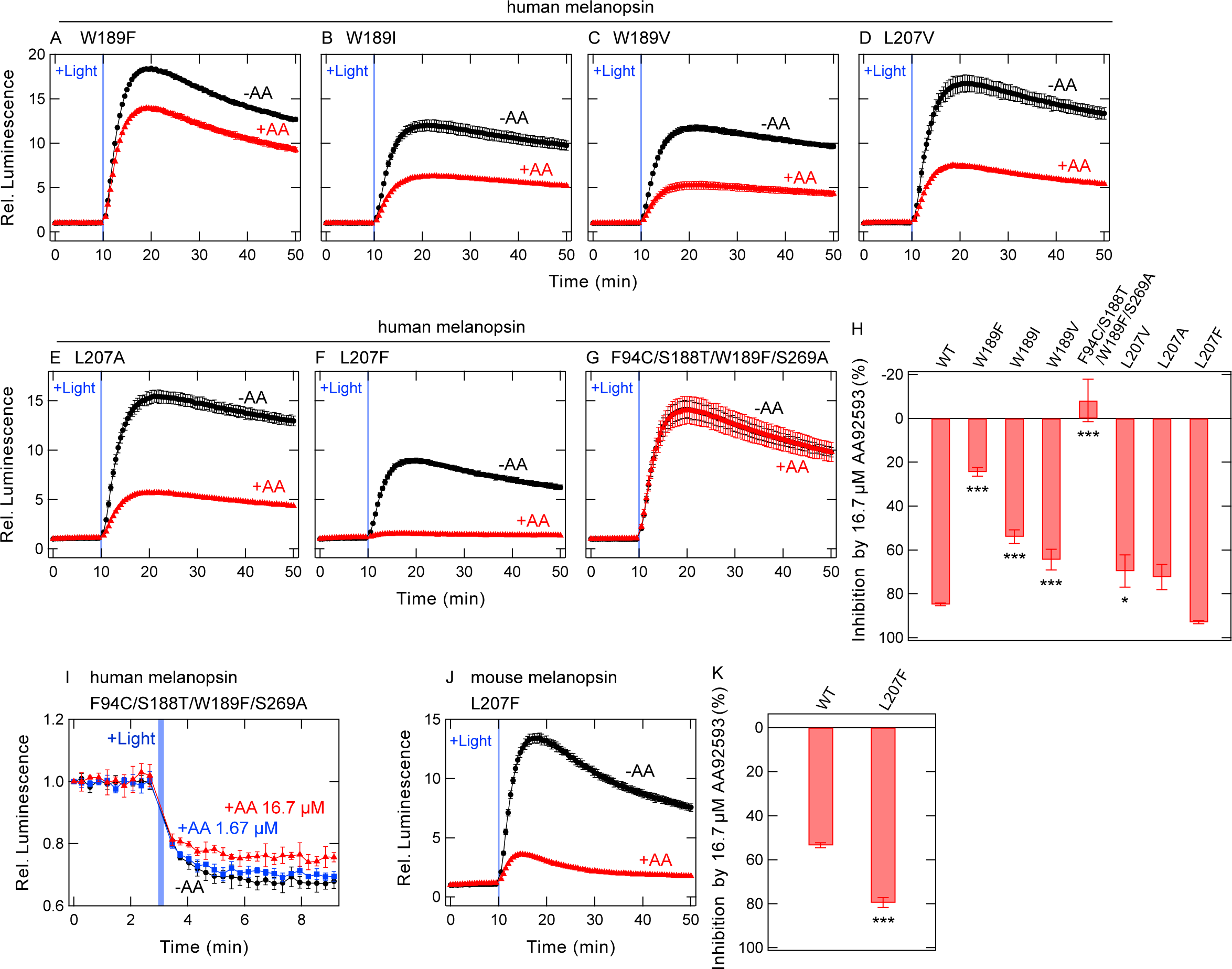
Changes in susceptibility of human melanopsin for AA92593 by substitutions of amino acid residues interacting with AA92593 in simulations. (*A*-*G*) AA92593-dependent inhibition of intracellular cAMP elevation in COS-1 cells upon Gsα/q11 activation of human melanopsin mutants W189F (*A*), W189I (*B*), W189V (*C*), L207V (*D*), L207A (*E*), L207F (*F*), and F94C/S188T/W189F/S269A (*G*). Error bars indicate the SD values (n = 3). (*H*) Comparison of inhibition in peak cAMP responses by 16.7 μM AA92593 upon Gsα/q11 activation in human melanopsin WT and mutants. WT data is the same as Fig. 2G. Error bars indicate the SD values (n = 46, 3, 3, 3, 3, 3, 3, and 3 for WT, W189F, W189I, W189V, F94C/S188T/W189F/S269A, L207V, L207A, and L207F, respectively). The statistical *P* values in differences from WT are <0.001***, <0.001***, <0.001***, <0.001***, 0.010**, 0.07, and 0.59 for W189F, W189I, W189V, F94C/S188T/W189F/S269A, L207V, L207A, and L207F, respectively (Dunnett’s test following one-way ANOVA). (*I*) NanoBiT Gq dissociation assay using Gqα/R183Q-LgBiT on human melanopsin F94C/S188T/W189F/S269A mutant. Red, blue, and black traces indicate luminescence changes in the presence of 16.7 μM (red), 1.67 μM (blue), and no (black) AA92593. Light blue bars indicate white light illumination (10 sec). NanoLuc luminescence levels are normalized to the values at the starting point (time = 0 min). Error bars indicate the SD values (n = 3). (*J*) AA92593-dependent inhibition of intracellular cAMP elevation in COS-1 cells upon Gsα/q11 activation of mouse melanopsin L207F mutant. Error bars indicate the SD values (n = 3). In panels *A*-*G*, and *J*, Red and black traces indicate luminescence changes in the presence and absence of 16.7 μM AA92593, respectively. Light blue bars indicate white light illumination (10 sec). Luminescence levels of cAMP biosensor (GloSensor) are normalized to the values at the starting point (time = 0 min). (*K*) Comparison of inhibition in peak cAMP responses by 16.7 μM AA92593 upon Gsα/q11 activation in mouse melanopsin WT and L207F mutant. WT data is the same as Fig. 3E. Error bar indicates the SD value (n = 23 and 3 for WT and L207F, respectively). The statistical *P* values of L207F in difference from WT is <0.001*** (Dunnett’s test following one-way ANOVA).

**Fig. 6.**
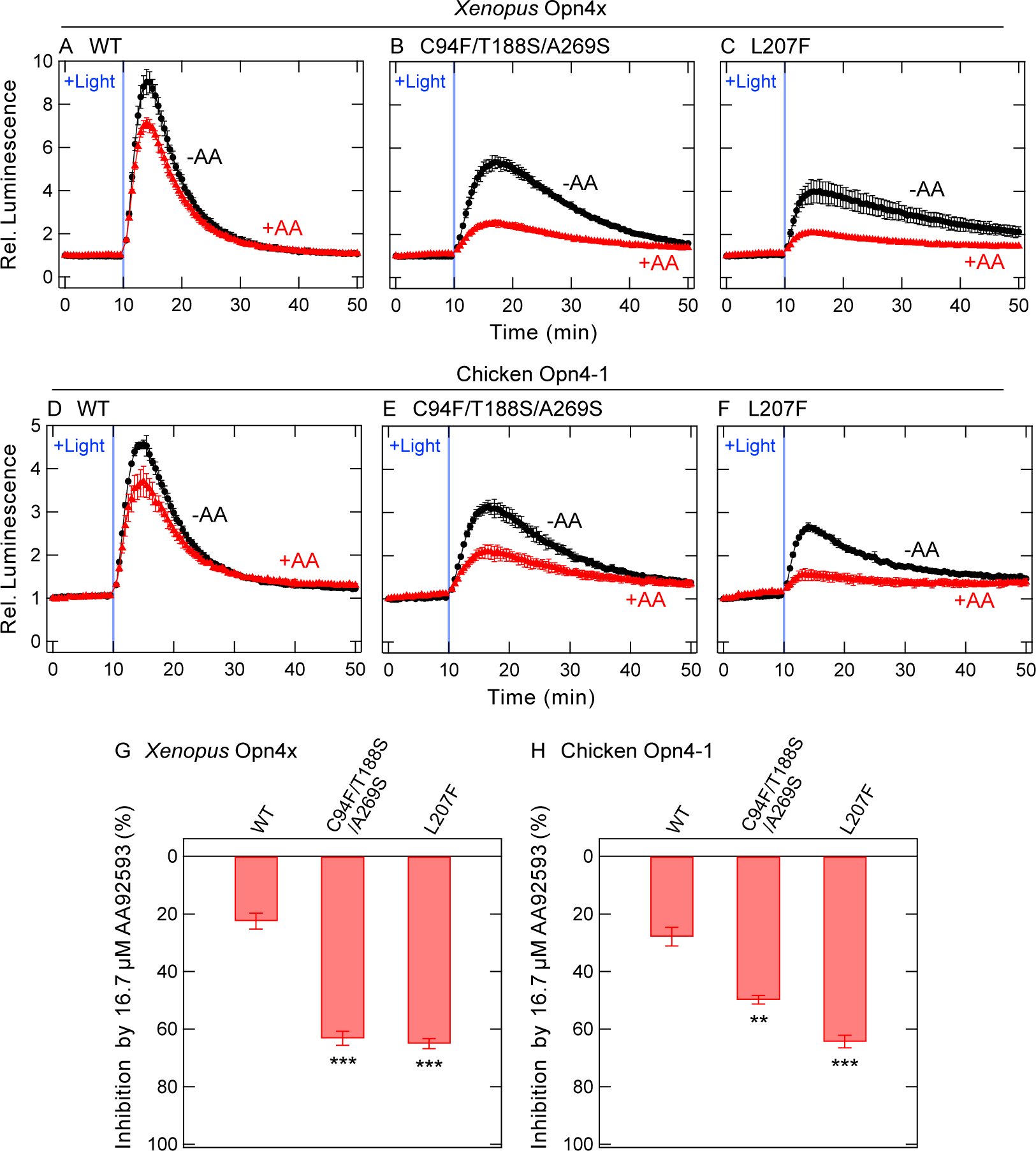
Changes in susceptibility of non-mammalian-type melanopsins for AA92593 by amino acid substitutions. (*A*-*C*) AA92593-dependent inhibition of intracellular cAMP elevation in COS-1 cells upon Gsα/q11 activation of *Xenopus* Opn4x WT (*A*) as well as mutants C94F/T188S/A269S (*B*) and L207F (*C*). (*D*-*F*) AA92593-dependent inhibition of intracellular cAMP elevation in COS-1 cells upon Gsα/q11 activation of chicken Opn4-1 WT (*D*) as well as mutants C94F/T188S/A269S (*E*) and L207F (*F*). In panels *A*-*F*, red and black traces indicate luminescence changes in the presence and absence of 16.7 μM AA92593, respectively. Light blue bars indicate white light illumination (10 sec). Luminescence levels of cAMP biosensor (GloSensor) are normalized to the values at the starting point (time = 0 min). Error bars indicate the SD values (n = 3). (*G* and *H*) Comparison of inhibition in peak cAMP responses by 16.7 μM AA92593 upon Gsα/q11 activation in WT and mutant of *Xenopus* Opn4x (*G*) and chicken Opn4-1 (*H*). Error bars indicate the SD values (n = 8, 4, and 3 for *Xenopus* Opn4x WT, C94F/T188S/A269S, and L207F, respectively, and 8, 3, and 3 for chicken Opn4-1 WT, C94F/T188S/A269S, and L207F, respectively). The statistical *P* values in differences from WT are <0.001*** and <0.001*** for *Xenopus* Opn4x C94F/T188S/A269S and L207F, respectively, and 0.003** and <0.001*** for chicken Opn4-1 C94F/T188S/A269S and L207F, respectively (Dunnett’s test following one-way ANOVA).

### Experimental assessment of interaction of Trp-189 and Leu-207 with AA92593

Based on the docking and MD simulations of human melanopsin with AA92593, importance of interactions of Trp-189 and Leu-207 with the antagonist was experimentally examined using the GsX GloSensor assay. To validate the importance of Trp-189 in the interaction with AA92593, substitutions of W189F, W189I, or W189V in human melanopsin were tested. The inhibitory effect of AA92593 was significantly weaker in all the three Trp-189 mutants than in WT (Fig. 5, A, B, C, and H). In particular, the W189F substitution resulted in a more marked decrease in sensitivity to AA92593 than other single substitutions (Fig. 5A). To assess the importance of Leu-207 in the interaction with AA92593, substitutions L207V, L207A, or L207F in human melanopsin were tested. The L207V and L207A mutants showed significantly reduced antagonistic effects of AA92593, similar to other substitutions at the AA92593 binding site (Fig. 5, D, E, and H). Of note, the L207F substitution enhanced the sensitivity of human melanopsin to AA92593 (Fig. 5F), although the difference was not statistically significant (Fig. 5H). Thus, we tested the effect of L207F substitution on mouse melanopsin, which is less sensitive to AA92593 (Fig. 1B). As expected, the mouse melanopsin L207F mutant showed significantly increased susceptibility to the antagonist (Fig. 5, J and K). The introduction of a Phe residue at position 207 caused additional interactions between the phenyl ring in Phe-207 and some functional groups in AA92593. The marked changes in the antagonistic effects of Trp-189 and Leu-207 substitutions are consistent with our molecular simulations, showing that these residues are located at the antagonist-binding site (Fig. 4B).

As mentioned above, the W189F mutant showed the lowest sensitivity to AA92593 among the analyzed single mutants (Fig. 5, A and H). Next, we attempted to render human melanopsin more ineffective to AA92593 by combining W189F with F94C/S188T/S269A. The quadruple F94C/S188T/W189F/S269A mutant of human melanopsin almost completely lost sensitivity to AA92593 (Fig. 5, G and H). The NanoBiT G protein dissociation assay of the F94C/S188T/W189F/S269A mutant also showed little sensitivity to AA92593 (Fig. 5I), although a synergistic effect of W189F and F94C/S188T/S269A was not clearly observed (Fig. 5G). This is because, in the NanoBiT assay, the triple mutant F94C/S188T/S269A (without W189F) was already almost insensitive to AA92593 (Fig. 2I). In the GloSensor assay, a small difference in G protein activation between the triple and quadruple mutants in the presence of AA92593 would be amplified through the second messenger signaling cascade.

Taken together data from GloSensor and NanoBiT assays combined with MD simulations, we concluded that amino acid residues at positions 94, 188, 269, 189, and 207 in mammalian melanopsins presumably form the antagonist-binding site and play critical roles in susceptibility to AA92593. Thus, the interaction with AA92593 can be regulated by these amino acids. Next, we tested whether we could make non-mammalian melanopsin sensitive to the antagonist by substitutions at these sites.

### Inducing the susceptibility of non-mammalian-type melanopsin (Opn4x) to AA92593 by amino acid substitutions

Our results showed that Phe-94, Ser-188, Ser-269, Trp-189, and Leu-207 were important for specific interactions of mammalian melanopsin with AA92593; therefore, we attempted to make non-mammalian-type melanopsins (also known as Opn4x) susceptible to the antagonist by introducing amino acid substitution(s). We noticed *Xenopus* Opn4x (36) and chicken Opn4-1 (37, 38) as typical non-mammalian-type melanopsins, and we introduced Phe-94/Ser-188/Ser-269 into them (see Fig. 2A). We expected that the “mammalian melanopsin-type” amino acid residues would increase sensitivity of the non-mammalian-type melanopsins for the antagonist. Trp-189 and Leu-207 are conserved in the non-mammalian-type melanopsins (Fig. S2). We did not substitute Trp-189, and introduced the L207F substitution because this substitution increased susceptibility of mammalian melanopsins to the antagonist (Fig. 5, F, H, J, and K).

The GsX GloSensor assay using Gsα/q11 of *Xenopus* Opn4x and chicken Opn4-1 detected light-dependent Gq activation by both non-mammalian-type melanopsins (Fig. 6, A and D). Unlike mammalian melanopsins (Fig. 1), the addition of AA92593 to *Xenopus* Opn4x and chicken Opn4-1 did not effectively inhibit the light-dependent responses (Fig. 6, A and D). These results indicated that AA92593 is not effective as an antagonist for the non-mammalian-type melanopsins, consistent with the fact that Phe-94, Ser-188, and Ser-269 are not conserved among non-mammalian-type melanopsins (Figs. 2A and S2). If these three residues are important for the antagonistic effect against melanopsins, the introduction of Phe-94/Ser-188/Ser-269 into the non-mammalian-type melanopsins would make them susceptible to AA92593. As expected, the substitutions of C94F/T188S/A269S in *Xenopus* Opn4x and chicken Opn4-1 increased their sensitivity to AA92593 (Fig. 6, B, E, G, and H). Based on the loss-of-function properties of F94C/S188T/S269A mutants of mammalian-type melanopsins and the gain-of-function properties of C94F/T188S/A269S mutants of non-mammalian-type melanopsins (Figs. 2G, 3E, 6G, and 6H), we propose that Phe-94, Ser-188, and Ser-269 are required for specific interactions with AA92593 as an effective antagonist.

Similar to human and mouse melanopsins (Fig. 5, F, H, J, and K), the introduction of a Phe residue at position 207 in *Xenopus* Opn4x and Chicken Opn4-1 increased their susceptibility of the melanopsins for AA92593, although the substitutions reduced light-dependent cAMP responses by the melanopsins (Fig. 6, C, F, G, and H). The results of substitutions at position 207 supported the idea that AA92593 binds to a similar site in both non-mammalian-type and mammalian-type melanopsins, and substitutions at positions 94, 188, 269, and 207 make the non-mammalian-type melanopsins interact with the antagonist in a similar way to AA92593-sensitive mammalian-type melanopsins.

### Inducing the susceptibility of invertebrate melanopsin to AA92593 by amino acid substitutions

Non-mammalian-type melanopsins were successfully converted into AA92593-susceptible types by introducing Phe-94/Ser-188/Ser-269 or Phe-207 (Fig. 6). We attempted to extend the functional conversion targets to invertebrate melanopsin. Cephalochordate amphioxus species *Branchiostoma belcheri* and *Branchiostoma lanceolatum* possess melanopsin (15, 39), and in this study, we indicate these melanopsins as *belcheri* and *lanceolatum* melanopsins, respectively. We assessed the susceptibility of *belcheri* and *lanceolatum* melanopsins to AA92593 using the GsX GloSensor assay with Gsα/q11. In the absence of AA92593, the *belcheri* melanopsin produced smaller light-dependent cAMP responses than the other melanopsins assessed in this study (Fig. 7A), whereas the *lanceolatum* melanopsin produced robust cAMP responses upon light illumination (Fig. 7B). Surprisingly, light-induced cellular responses of the *belcheri* melanopsin were reduced by approximately 40% by the addition of AA92593 (Fig. 7A). On the other hand, responses by the *lanceolatum* melanopsin was insensitive to the antagonist (Fig. 7B), similar to non-mammalian-type melanopsins (Fig. 6, A and D). Although the mechanism underlying the AA92593 susceptibility of *belcheri* melanopsin is unknown, we attempted to convert AA92593-insensivite *lanceolatum* melanopsin to a susceptible type via amino acid substitutions.

**Fig. 7.**
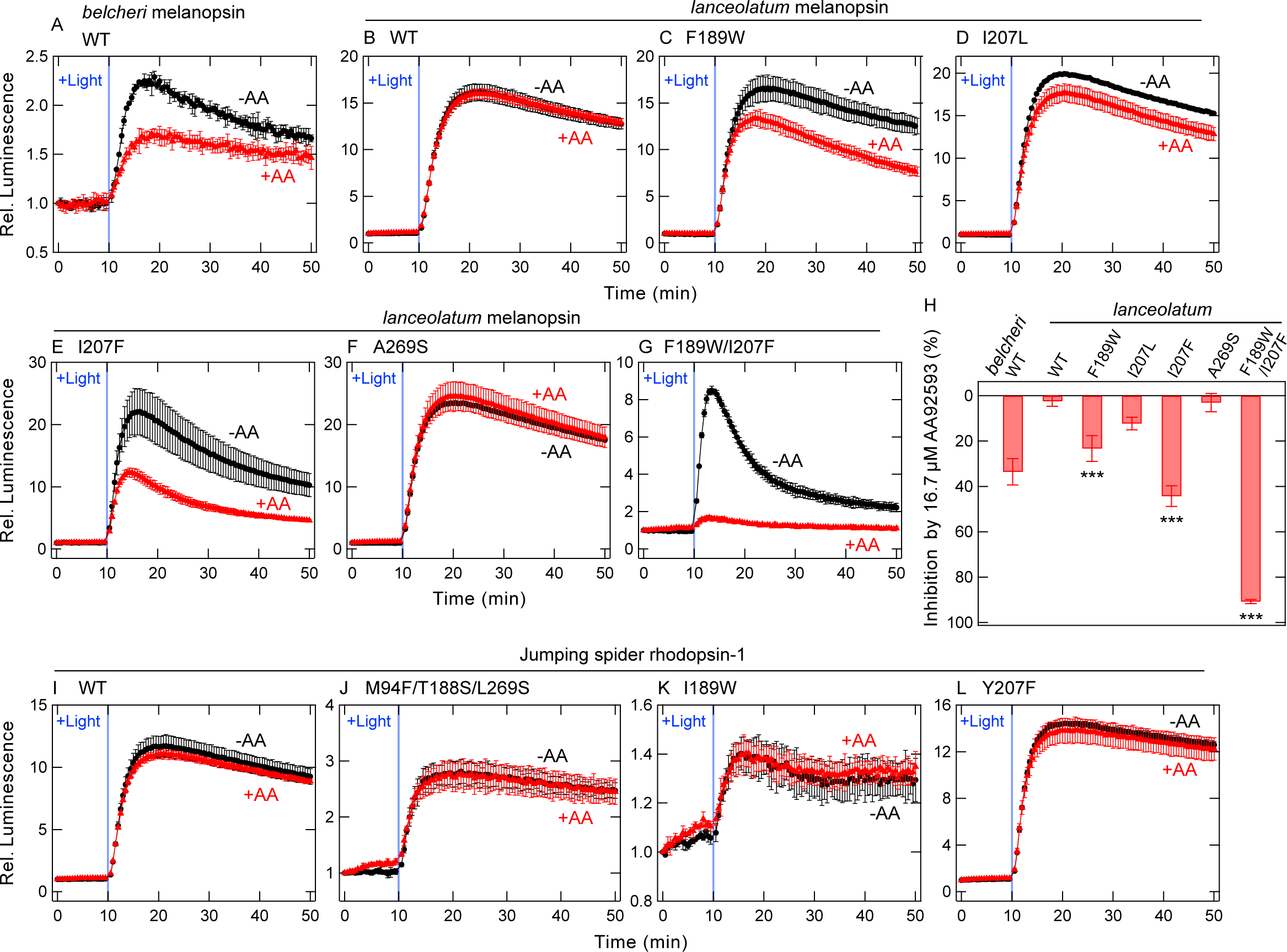
Changes in susceptibility of invertebrate melanopsins and a non-melanopsin Gq-coupled opsin for AA92593 by amino acid substitutions. (A) AA92593-dependent inhibition of intracellular cAMP elevation in COS-1 cells upon Gsα/q11 activation of *belcheri* melanopsin WT. Error bars indicate the SD values (n = 3). (*B*-*G*) AA92593-dependent inhibition of intracellular cAMP elevation in COS-1 cells upon Gsα/q11 activation of *lanceolatum* melanopsin WT (*B*) as well as mutants F189W (*C*), I207L (*D*), I207F (*E*), A269S (*F*), and F189W/I207F (*G*). Error bars indicate the SD values (n = 3). (*H*) Comparison of inhibition in peak cAMP responses by 16.7 μM AA92593 upon Gsα/q11 activation in *belcheri* melanopsin WT and *lanceolatum* melanopsin WT and mutants. Error bars indicate the SD values (n=3, 8, 4, 3, 3, 3, and 3 for *belcheri* WT, *lanceolatum* WT, F189W, I207L, I207F, A269S, F189W/I207F, respectively). The statistical *P* values in differences from WT are <0.001***, 0.23, <0.001***, 1.00, and <0.001*** for F189W, I207L, I207F, A269S, F189W/I207F, respectively (Dunnett’s test following one-way ANOVA). (*I*-*L*) AA92593-dependent inhibition of intracellular cAMP elevation in COS-1 cells upon Gsα/q11 activation of jumping spider rhodopsin-1 WT (*I*), M94F/T188S/L269S (*J*), I189W (*K*), and Y207F (*L*). Error bars indicate the SD values (n = 3). In panels *A*-*G* and *I*-*L*, Red and black traces indicate luminescence changes in the presence and absence of 16.7 μM AA92593, respectively. Light blue bars indicate white light illumination (10 sec). Luminescence levels of cAMP biosensor (GloSensor) are normalized to the values at the starting point (time = 0 min).

Because *lanceolatum* melanopsin endogenously possesses Phe-94 and Ser-188 (Fig. 2A), we substituted Ala-269 with Ser, Phe-189 with Trp, and Ile-207 with Leu or Phe in the invertebrate melanopsin. We assessed effect of the antagonist on the mutants using the GsX GloSensor assay with Gsα/q11. F189W and I207F substitutions significantly increased the sensitivity to AA92593 (Fig. 7, C, E, and H), whereas I207L and A269S caused little changes in sensitivity (Fig. 7, D, F, and H). Furthermore, the double substitution F189W/I207F further increased the susceptibility of the *lanceolatum* melanopsin to the antagonist (Fig. 7G). The AA92593-induced inhibition of the double mutant F189W/I207F was as efficient as that of human melanopsin (Fig. 1A). Taken together, the introduction of Trp-189 and Phe-207 rendered *lanceolatum* melanopsin highly sensitive to the antagonist (Fig. 7H).

We next tested the effect of AA92593 on jumping spider rhodopsin-1, a Gq-coupled visual opsin but not melanopsin (see Figs. 2 and S2) (40). Similar to non-mammalian-type and invertebrate melanopsins, jumping spider rhodopsin-1 was not sensitive to AA92593 as assessed using the GsX GloSensor assay (Fig. 7I). The introduction of Phe-94/Ser-188/Ser-269, Trp-189, or Phe-207 into jumping spider rhodopsin-1 did not increase its sensitivity to AA92593 (Fig. 7, J, K, and L), unlike in the cases of non-mammalian-type and invertebrate melanopsins (Figs. 6 and 7). The introduction of I189W/Y207F impaired the light-dependent cAMP responses even in the absence of AA92593 (Fig. S8). Taken together, the non-melanopsin Gq-coupled opsin cannot be functionally converted to an AA92593-sensitive type by site-directed mutagenesis, in contrast to non-mammalian-type and invertebrate melanopsins.

## Discussion

In the present study, we investigated the molecular mechanisms underlying the antagonistic effects of AA92593 on mammalian melanopsins. A combination of cell-based assays and computational simulations identified five amino acid residues comprising the antagonist-binding site and responsible for the antagonism. Based on these results, we discuss how AA92593 acts as the antagonist specifically against mammalian melanopsins. We propose that the insights gained from this study can be applied to further physiological studies.

### Molecular basis underlying the specific interaction between mammalian melanopsin and AA92593

Our site-directed mutagenesis data showed that amino acid substitutions at positions 94, 188, 269, 189, and 207 decreased or increased the antagonistic effects of AA92593 (Figs. 2, 3, and 5). Our docking and MD simulations of human melanopsin revealed that these amino acids are located at the AA92593-binding site (Fig. 4). The most effective single substitution site reducing the effect of AA92593 on mammalian melanopsin was Trp-189, which is conserved among mammalian-type and non-mammalian-type melanopsins (see Fig. S2). In the simulated structural models of human melanopsin (Fig. 4), Trp-189 was found to interact with the antagonist from the extracellular side, primarily via van der Waals interactions (Figs. 4B and S6). These consistent experimental and simulation results suggest that these five amino acid residues modulate the antagonistic effects through direct interactions. This interpretation is supported by the fact that the introduction of “mammalian melanopsin-type” amino acid residues (Phe-94, Ser-188, and Ser-269) as well as Trp-189 and Phe-207 into non-mammalian and invertebrate melanopsins made them more susceptible to the antagonist (Figs. 6 and 7).

Substitutions at the five sites converted AA92593-insensitive melanopsins to sensitive ones (Figs. 6 and 7), but did not convert jumping spider rhodopsin-1, a non-melanopsin Gq-coupled opsin (Fig. 7, I-L). These results suggest that the molecular architecture around the AA92593-binding site is somewhat different between melanopsins and other Gq-coupled opsins. The differences in the molecular architecture between melanopsins and other opsins would be useful for designing chemicals that selectively bind to either melanopsins or other opsins.

In addition to the five amino acid residues, Trp-265 was also found to be in contact with the antagonist from the intracellular side and showed strong binding. Because Trp-265 is highly conserved in GPCRs and is expected to be important for their function (32, 35, 41), this strong interaction between AA92593 and Trp-265 may play a role in suppressing the activation of human melanopsin.

### Modulation of AA92593 sensitivity in melanopsins for physiological studies

Our data clearly indicated that the susceptibility to AA92593 as an antagonist can be modulated by substituting amino acid residues in a wide variety of melanopsins. Substitutions at positions 94/188/269/189/207 decreased or increased the susceptibility of mammalian and other melanopsins. Most mammals, including humans and mice, possess a single melanopsin gene (opn4m) in their genomes (38), and Opn4m primarily functions in ipRGCs. In contrast, many non-mammalian vertebrates possess multiple melanopsin genes (opn4m and opn4x). Melanopsins of non-mammalian vertebrates are expressed in various cells, including ipRGCs, but the physiological role of each melanopsin remains unclear (38, 42, 43). Our results showed that AA92593 is more effective against mammalian-type melanopsin (Opn4m) rather than against non-mammalian-type one (Opn4x), and that the effectiveness of AA92593 on a specific melanopsin can be more accurately predicted based on the amino acid sequence at the five sites (Fig. S2). In addition, susceptibility of invertebrate melanopsins to AA92593 was dramatically changed by the introduction of Trp-189 and Phe-207 (Fig. 7G).

Recent progress in genome editing techniques (44) has enabled the elimination of a specific melanopsin gene in animals, but temporal silencing of specific melanopsin functions remains difficult. Genome editing to introduce amino acid substitutions at positions 94/188/269/189/207 into a specific melanopsin can increase or decrease the susceptibility of the target melanopsin to AA92593. In particular, the introductions of Trp-189 or Phe-207 were the most effective single substitutions to increase AA92593 antagonism (Figs. 6 and 7). Such a gene manipulation to introduce amino acid residue(s) that modulate the effect of AA92593 will enable the antagonist-mediated suppression of the targeted melanopsin functions. The combination of site-directed genome editing and AA92593 administration in living animals could silence specific melanopsin functions with high spatiotemporal resolution.

## Experimental Procedures

### Constructs

In the constructs for melanopsin expression (opsin name - GenBank ID: human melanopsin - NM_033282.4, mouse melanopsin - NM_013887.2, *Xenopus* Opn4x - NM_001085674.1, chicken opn4-1 - NM_001397961.1, *Branchiostoma belcheri* melanopsin - AB205400.1, and *Branchiostoma lanceolatum* melanopsin - MF464477.1), the C-terminal amino acid residues (94, 140, 200, 214, 296, and 308 residues for human (9), mouse (9), *Xenopus*, chicken, *belcheri* (15), and *lanceolatum* melanopsins, respectively) were removed, the 1D4 tag sequence (ETSQVAPA) was added, and inserted into the EcoRI/NotI site in a mammalian expression vector pMT. Jumping spider rhodopsin-1 (GenBank ID: AB251846.1) was also added with the 1D4 tag, and inserted into pMT vector. Human and mouse melanopsins, and jumping spider rhodopsin-1 in pCDNA3.1 were kindly provided from Drs. Akihisa Terakita and Mitsumasa Koyanagi (Osaka Metropolitan University). In this paper, each opsin with the C-terminal sequence modifications as described above is denoted “WT”. Bovine Gsα and its mutants Gsα/q11 and Gsα/i11 (C-terminal 11 residues are replaced with Gqα and Giα, respectively) were also inserted into the pMT vector. The expression vector pGlo-22F for GloSensor assay was purchased from Promega (Madison, WI). For NanoBiT assay, the cDNAs of Lg-BiT inserted human Gqα, human Gβ1, and the Sm-BiT fused human Gγ2 (C68S), RIC8A were constructed according to refs. (22) and (23), and inserted into the pMT vector. The expression vector for RIC8A was purchased from GenScript (Piscataway, NJ), and subcloned into the pMT vector.

To construct the various melanopsin mutants, Gsα/q11, Gsα/i11, and Gqα/R183Q-LgBiT, mutations were introduced into the cDNA sequence by PCR reaction. The sequences were confirmed by DNA sequencing. The plasmid DNA used for transfection was prepared using either the FastGene Plasmid Mini Kit (Nippon Genetics, Japan) or NucleoBond Xtra Midi/Maxi (TAKARA, Japan).

### Transfection to COS-1 cells for GloSensor assay and NanoBiT G protein dissociation assay

Opsins, GloSensor assay sensor (coded by pGlo-22F), and NanoBiT-tagged G proteins were transiently expressed in COS-1 cells (kindly provided from Dr. David Farrens, Oregon Health and Science University) using polyethyleneimine as described previously (23, 45). For the GsX GloSensor assay (12, 13), each well of a 96-well assay plate (Corning, Kennebunk, ME) was transfected with 50 ng opsin plasmid, 16.7 ng G protein plasmid, 50 ng pGlo-22F plasmid (Promega), and 500 ng polyethyleneimine in 25 μL Opti-mem (Gibco, Waltham, MA) and 75 μL D-MEM (Wako, Japan) containing 10 % (v/v) FBS, 100 units/mL penicillin, and 100 μg/mL streptomycin. For NanoBiT G protein dissociation assay, each well of a 96-well assay plate was transfected with 50 ng opsin plasmid, 5 ng Lg-BiT inserted Gqα (Gqα-LgBiT) plasmid, 25 ng Gβ1 plasmid, 25 ng Sm-BiT fused Gγ2 plasmid, 5 ng RIC8A plasmid, and 500 ng polyethyleneimine in 25 μL Opti-mem (Gibco) and 75 μL D-MEM (Wako) containing 10% (vol/vol) FBS, 100 units/mL penicillin, and 100 μg/mL streptomycin.

### GsX GloSensor assay

The transfected COS-1 cells were incubated at 37°C, 5% CO_2_ for 2 days, and the medium was aspirated and exchanged with 50 μL of HBSS (145 mM NaCl, 10 mM D-glucose, 5 mM KCl, 1 mM MgCl_2_, 1.7 mM CaCl_2_, 1.5 mM NaHCO_3_, 10 mM HEPES, pH 7.4), containing 33.3, 20, 10, 2, 0.2, 0.02, or 0 μM AA92593, 0.5% DMSO, and 4% (vol/vol) GloSensor cAMP reagent stock solution. Under our experimental conditions, >50 μM AA92593 was not fully dissolved in the HBSS solution, and we set maximal concentration of AA92593 (Glixx Laboratories, Hopkinton, MA) at 33.3 μM (final concentration of 16.7 μM, see below). Then, the cells were incubated at room temperature for 1 h, and luminescence changes were monitored. After incubation of the cells with AA92593, luminescence measurement was interrupted, the plate was ejected, and the cells were added with 50 μL of HBSS containing 0.2 μM 9-*cis*-retinal. The final exogenous AA92593 concentrations were 16.7, 10, 5, 1, 0.1, 0.01, or 0 μM, and retinal concentration was 0.1 μM. 20 mM stock solution of AA92593 was prepared in DMSO, and final concentration of DMSO was kept at 0.25% regardless final AA922593 concentration. Then, the cells were incubated at room temperature for 1 h. Luminescence was measured using GM-2000 or GM-3510 microplate reader (Promega). Luminescence level of each well was measured every 30 s with integration time of 0.6-0.9 s. To illuminate opsins, luminescence measurement was interrupted, the plate was ejected, and the plate was illuminated by white light with CN-160 LED video light (light intensity, ∼9 mW/cm^2^) (NEEWER, China). After illumination, luminescence measurement was resumed. The measured luminescence levels were normalized to the level at the starting point (time = 0 min).

### NanoBiT G protein dissociation assay

The transfected COS-1 cells were incubated at 37°C, 5% CO_2_ for 1 day, and the medium was aspirated, followed by addition of 50 μL of HBSS containing 33.3 μM or 3.33 μM AA92593, 0.5% DMSO, and 10 μM coelenterazine h (Wako, Japan). Then, the cells were incubated at room temperature for 1 h. After incubation of the cells, luminescence measurement was interrupted, the plate was ejected, and the cells were added 50 μL of HBSS containing 0.2 μM 9-*cis*-retinal. The final concentration of AA92593 were 16.7 μM or 1.67 μM (DMSO concentration of 0.25%), and retinal concentration was 0.1 μM. Then, the cells were incubated at room temperature for 1 h. Luminescence was measured using GM-2000 or GM-3510 microplate reader (Promega). Luminescence level of each well was measured every 18 s with integration time of 0.3 s. To illuminate opsins, luminescence measurement was interrupted, the plate was ejected, and the plate was illuminated by white light with CN-160 LED video light (light intensity, ∼9 mW/cm^2^) (NEEWER, China). After illumination, luminescence measurement was resumed. The measured luminescence levels were normalized to the level at the starting point (time = 0 min).

### Docking and MD simulations

The predicted structure of the human melanopsin was taken from the AlphaFold Protein Structure Database (ID: Q9UHM6). The structure of AA92593 was built by GaussView (46). Docking calculations of the binding of AA92593 to the human melanopsin was performed using AutoDock Vina (25) and DiffDock (27). AutoDock Tools 1.5.4 (47) was used to set up the protein and ligand for AutoDock by deleting the water molecules and adding hydrogen atoms. Rigid-body docking was performed using AutoDock with the query box of 47 Å x 53 Å x 48 Å centered at (-14.5, 0.5, -0.1) to cover the expected binding pocket. The default parameters with inference_steps = 20, samples_per_complex = 40, and batch_size = 10 were used for DiffDock. The melanopsin-AA92593 complex with the best score from DiffDock was picked for the structural analysis and subsequent MD simulations. The initial structure for the simulation containing lipid and water molecules were prepared using the CHARMM-GUI Membrane Builder (48). The membrane was composed of 100% POPC with the KCl salt concentration of 150 mM, and the system size was 80.3 Å x 80.3 Å x 125.2 Å. The Amber FF14SB force field parameters were used for the proteins, and the waters were treated with the TIP3P water model. The long-range electrostatic interactions were treated by the particle mesh Ewald method, whereas short-range nonbonded interactions were cut off at 9 Å 10 Å during equilibration and production runs, respectively. Energy minimization with the steepest descent method was performed for 5000 steps. A series of short equilibrations were performed by gradually releasing the restraints on the position and dihedral of protein and membrane during a total of 1.875 ns by following the script generated by CHARMM-GUI. During these equilibration steps, the time step was increased from 1 to 2 fs and position restraints on the lipid headgroups were relaxed. Subsequently, all position and dihedral restraints were removed, and 10 ns simulation under the constant-NPT condition was performed to complete the equilibration. The Langevin thermostat and the Berendsen barostat were used to maintain the temperature at 303.15 K and pressure at 1 bar, respectively. A 1µs MD simulation under the constant-NPT condition at 303.15K and 1 bar was performed for production. To elucidate the key residues which have stronger influence on binding free energy, per-residue decompositions of the MMPBSA binding free energy were performed using AmberTools (49). The 1µs trajectory was used to calculate the binding free energy. Here, the negative decomposed binding free energy contribution indicates that the residue stabilizes binding and thus contributes to strengthen binding affinity. All molecular dynamics calculations were performed using Amber 22 software package (50).

## Author contributions

K. O., T. M., and H. T. designed the study. K. O. and H. T. conducted experiments, and analyzed obtained data. R. Z. and T. M. conducted docking and MD simulations and analyzed obtained data. All authors discussed experimental and simulational data and wrote the paper.

## Competing interests

The authors declare no competing interests.

## Abbreviations

ipRGC: intrinsically photosensitive retinal ganglion cell
GPCR: G protein-coupled receptor
MD: molecular dynamics
Opn4m: mammalian-type melanopsin
Opn4x: non-mammalian-type melanopsin
WT: wild-type

## Acknowledgment

We thank Dr. David Farrens (Oregon Health & Science University) for providing COS-1 cell line and the pMT expression vector, and Drs. Akihisa Terakita and Mitsumasa Koyanagi (Osaka Metropolitan University) for providing plasmids coding human and mouse melanopsins, and jumping spider rhodopsin-1. We also thank Hiroe Motomura and Kayo Inaba (Institute for Molecular Science), and the Functional Genomics Facility, NIBB Core Research Facilities (Okazaki, Japan) for technical support. The calculations were partially carried out at the Research Center for Computational Sciences in Okazaki (Project: 23-IMS-C111 and 24-IMS-C105).

## Funding

H. T. is supported by JST, PRESTO (JPMJPR1787) and the Japan Society for the Promotion of Science KAKENHI Grant 21H02445. T. M. is supported by the Japan Society for the Promotion of Science KAKENHI Grant 23K23303, 23KK0254, and 24K21756.

## Supporting Information

**Fig. S1.**
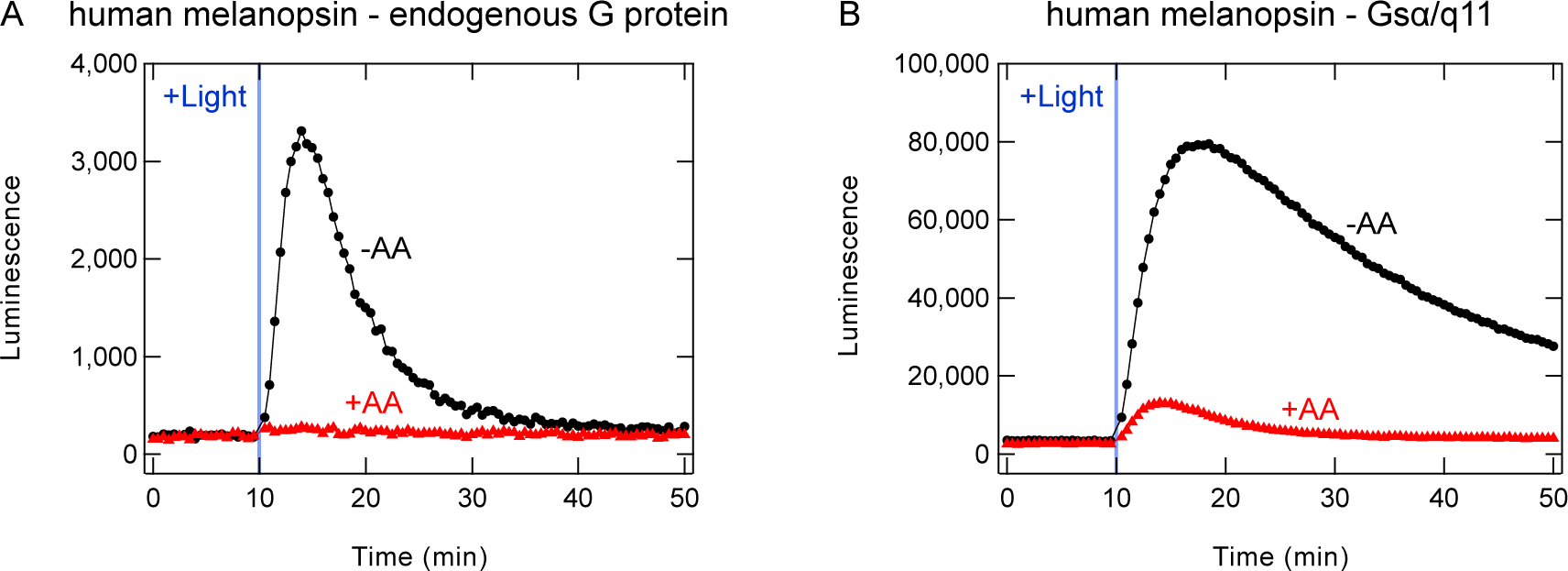
AA92593-dependent inhibition of intracellular cAMP elevation in COS-1 cells upon activation of endogenous G proteins and Gsα/q11 by human melanopsin. Representative raw luminescence changes upon activation of endogenous G proteins (*A*) and Gsα/q11 (*B*). Note that absolute luminescence level was ∼20-fold larger in Gsα/q11 activation, probably due to the high expression levels of the exogenous Gsα/q11 protein.

**Fig. S2.**
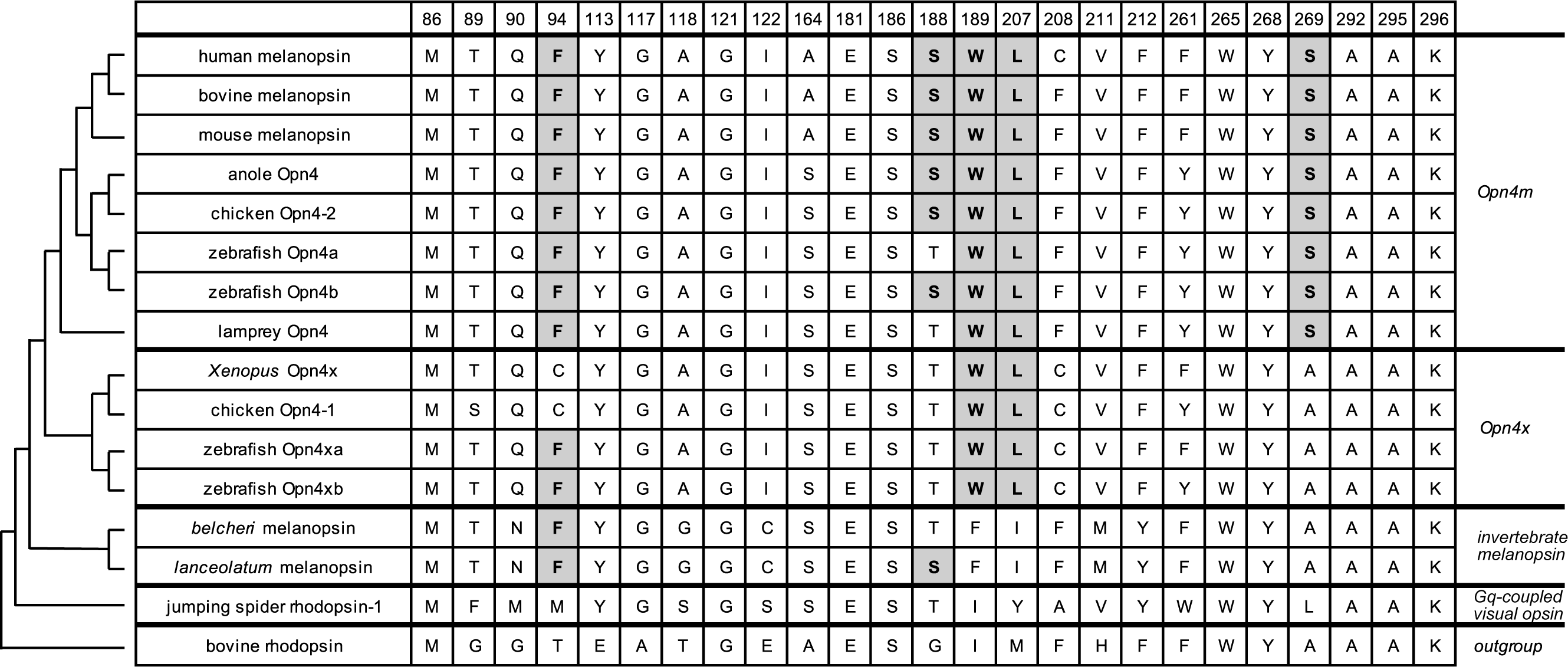
Comparison of amino acid residues in the retinal-binding site of various melanopsins and other opsins. Amino acid residues in the putative retinal-binding site of Opn4m, Opn4x, invertebrate melanopsin, non-melanopsin Gq-coupled opsin, bovine rhodopsin (amino acid numbering is based on bovine rhodopsin sequence) are shown, and schematic phylogenetic relationship of the opsins was also indicated.

**Fig. S3.**
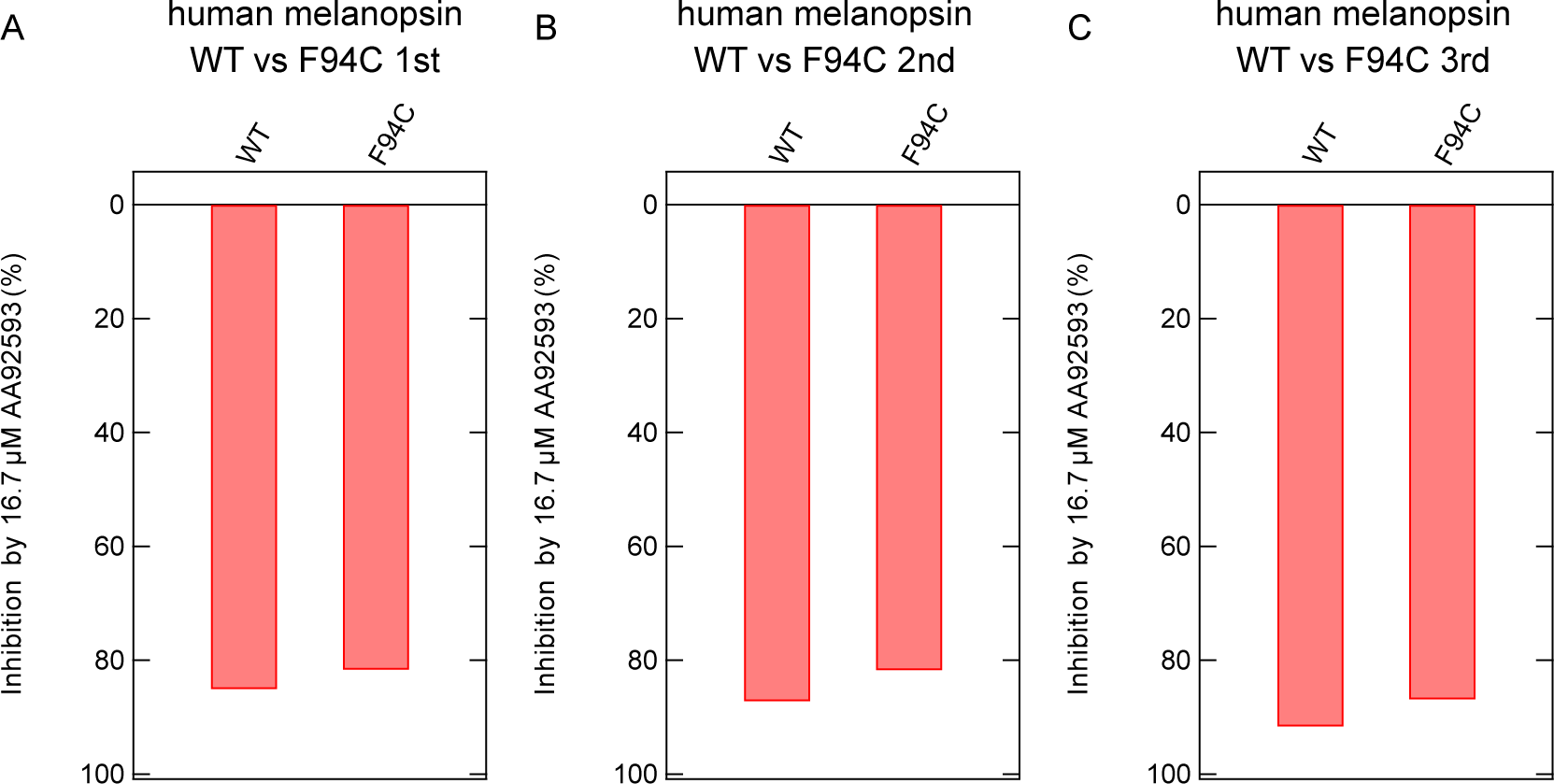
Comparison of inhibition in peak cAMP responses by 16.7 μM AA92593 upon Gsα/q11 activation in human melanopsin WT (*A*) and F94C mutant (*B*) in 3 times-repeated experiments. Note that in each experiment, F94C showed a less AA92593-induced suppression, although the difference was not statistically significant (see main text).

**Fig. S4.**
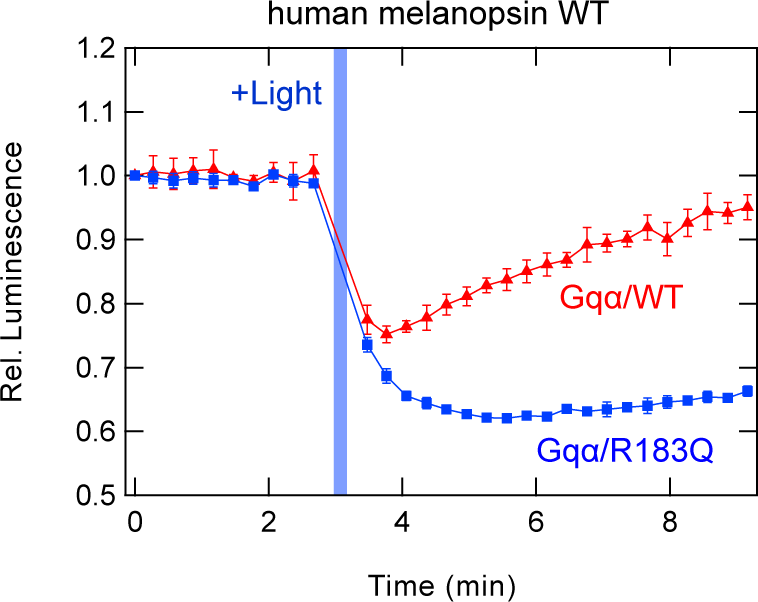
NanoBiT Gq dissociation assay using Gqα-LgBiT or Gqα/R183Q-LgBiT on human melanopsin WT. Red and blue traces indicate luminescence changes using Gqα-LgBiT and Gqα/R183Q-LgBiT, respectively, in the absence of AA92593. Light blue bars indicate white light illumination (10 sec). NanoLuc luminescence levels are normalized to the values at the starting point (time = 0 min). Error bars indicate the SD values (n = 3).

**Fig. S5.**
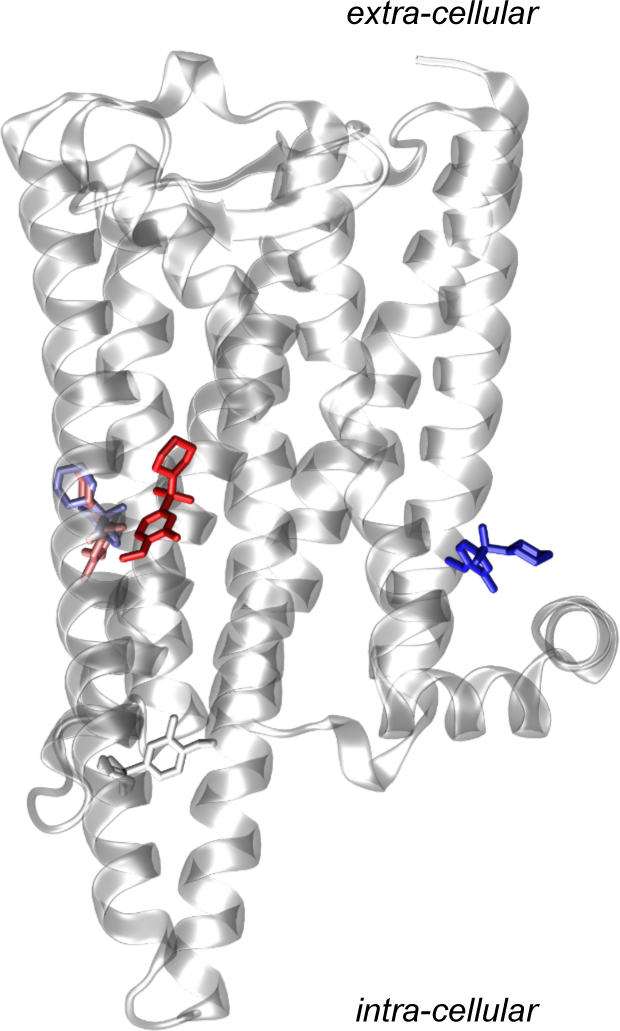
Docking structure of human melanopsin-AA92593 complex predicted using AutoDock. Location of predicted binding positions of AA92593 from AutoDock. Colors indicate different predicted positions.

**Fig. S6.**
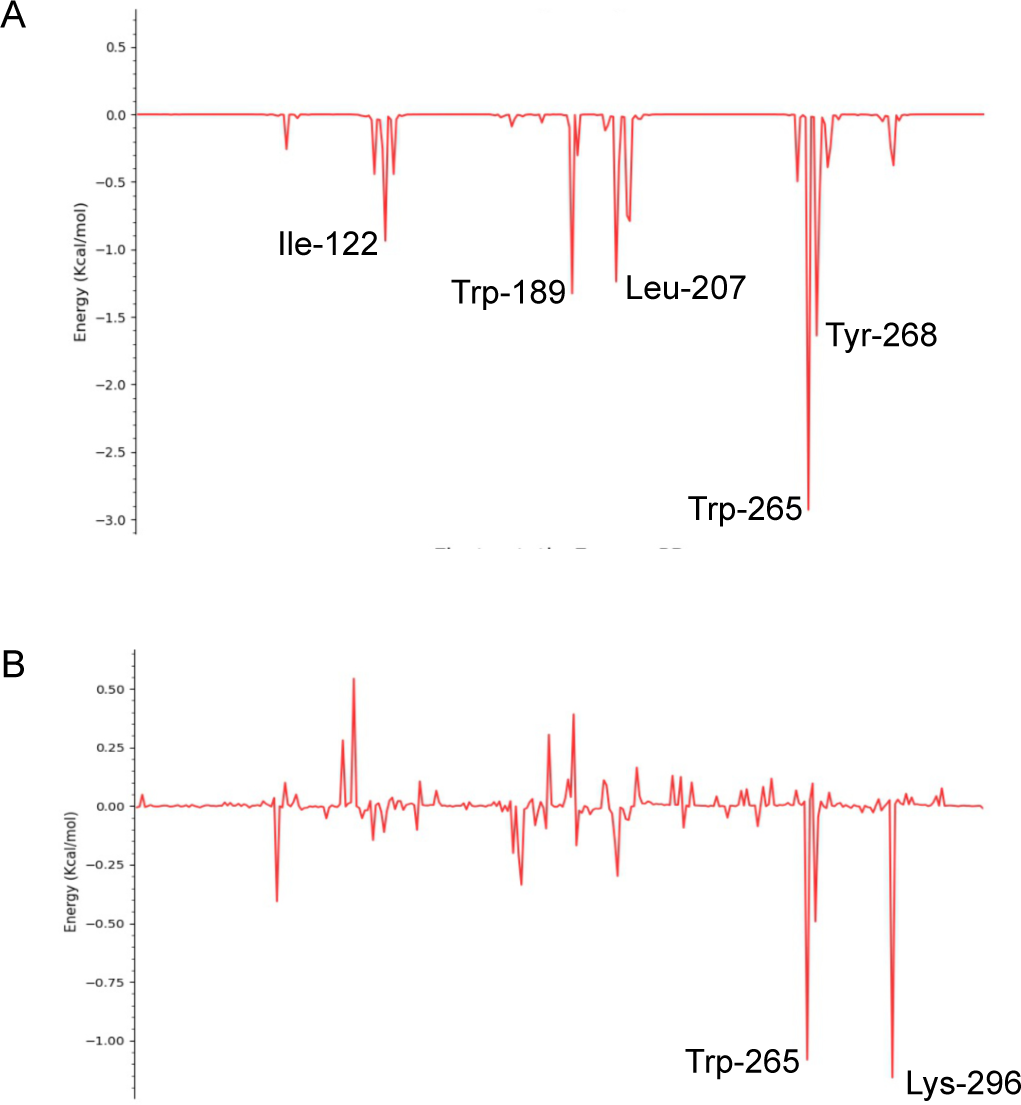
Energy decomposition of the per-residue decomposition for the binding of AA92593 to human melanopsin. Van der Waals (*A*) and electrostatic (*B*) contributions to the per-residue decomposition of the binding energy for the binding of AA92593 to human melanopsin calculated using MMPBSA.

**Fig. S7.**
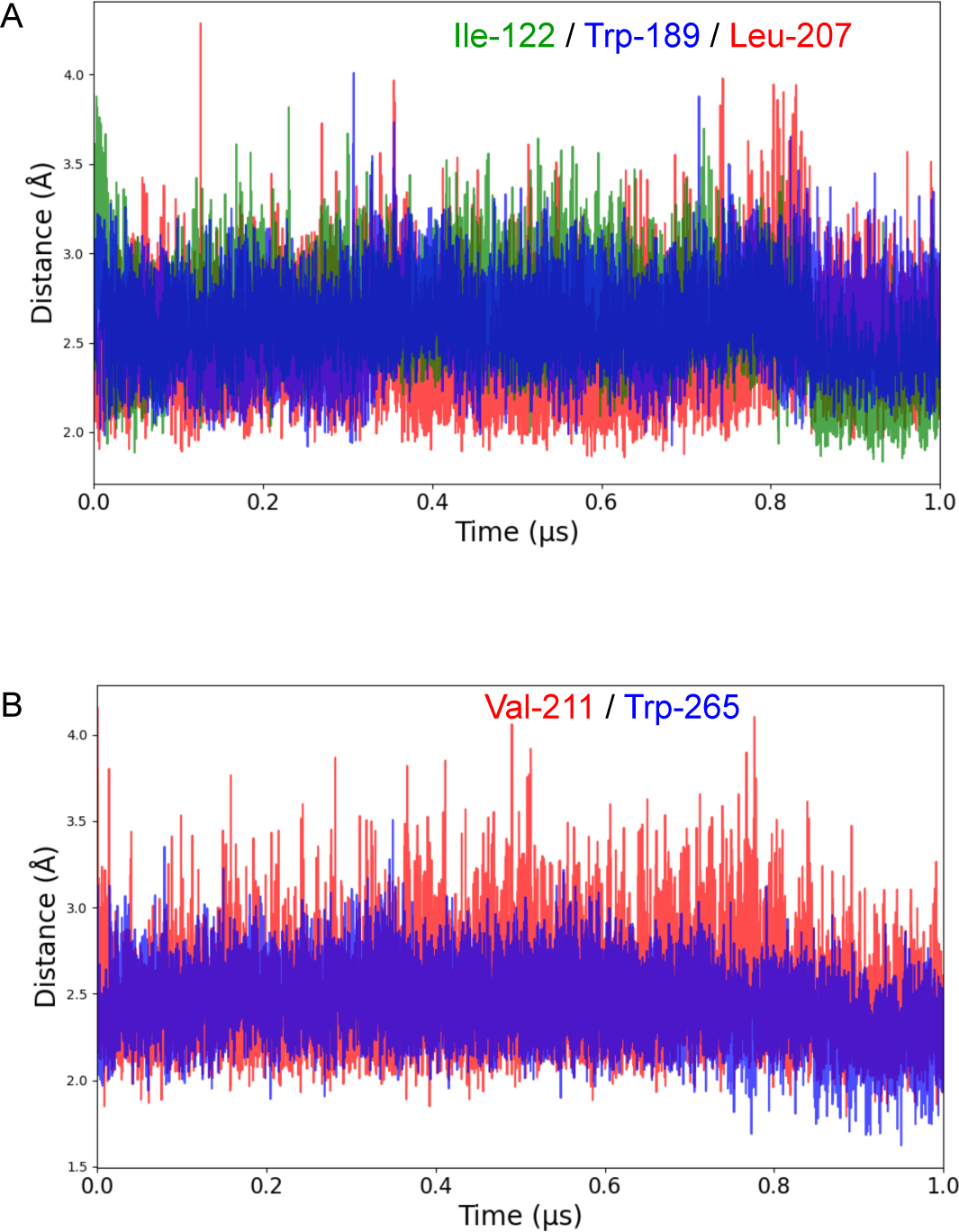
Changes of distances between AA92593 and selected residues in human melanopsin during MD simulation. Time evolution of the distances between AA92593 and selected residues in human melanopsin along the 1 µs MD trajectory of the complex. The results for Ile-122 (green), Leu-207 (red) and Trp-189 (blue) (in panel *A*), and Val-211 (red) and Trp-265 (blue) (in panel *B*) are shown.

**Fig S8.**
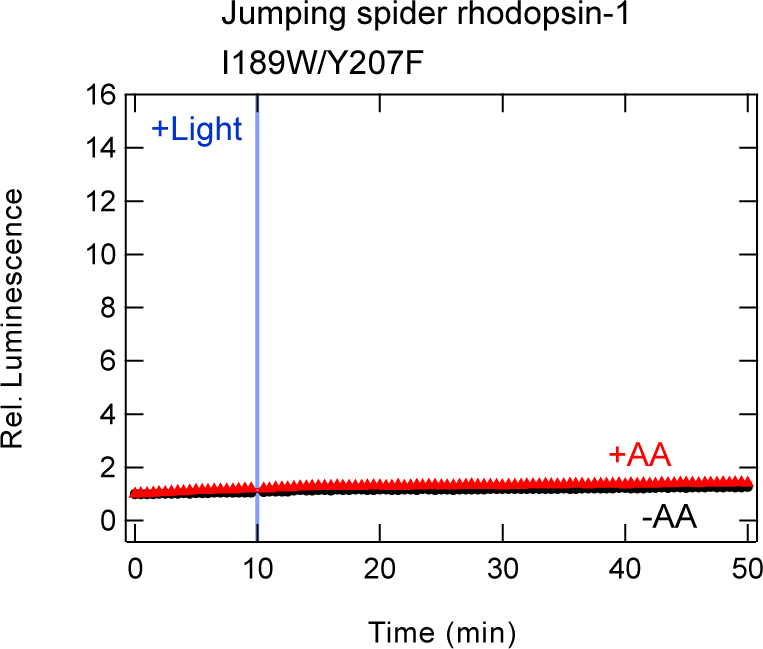
AA92593-dependent inhibition of intracellular cAMP elevation in COS-1 cells upon Gsα/q11 activation of jumping spider rhodopsin-1 I189W/Y207F mutant. Red and black traces indicate luminescence changes in the presence and absence of 16.7 μM AA92593, respectively. Light blue bars indicate white light illumination (10 sec). Luminescence levels of cAMP biosensor (GloSensor) are normalized to the values at the starting point (time = 0 min). Error bars indicate the SD values (n = 3).

## Notes

### Competing Interest Statement

The authors have declared no competing interest.

